# Isolation and genome sequencing of 14 *Spongia* sp. bacterial associates expands the taxonomic and functional breadth of the cultivatable marine sponge microbiome

**DOI:** 10.1101/2020.03.20.000216

**Authors:** Elham Karimi, Rodrigo Costa

**Affiliations:** Center of Marine Sciences-CCMAR, Faculty of Science and Technology, Campus of Gambelas, University of Algarve, 8005-139 Faro, Portugal; Institute for Bioengineering and Biosciences (IBB), Instituto Superior Tecnico, Universidade de Lisboa, 1049-001 Lisbon, Portugal; Department of Energy, Joint Genome Institute, Walnut Creek, California, USA, and Lawrence Berkeley National Laboratory, Berkeley, California, USA

**Keywords:** Porifera, genomics, host-microbe interactions, symbiosis, marine microbiology, secondary metabolite

## Abstract

Marine sponges live with complex microbial consortia, which have been considered as potential sources of novel natural products. However, the usual recalcitrance of host-associated microorganisms to cultivation makes studying sponge symbionts challenging. To tackle this complexity, exploration of cultivated sponge-associated bacteria and their coding potential is unavoidable. In this study, we isolate and report the draft genome sequences of 14 bacterial strains from the marine sponge *Spongia* sp. using R2A and VXA media. The strains belong to the classes *Actinobacteria, Gammaproteobacteria, Alphaproteobacteria,* and *Cytophagia* spanning 11 formally described genera plus two potentially novel genera in the *Rhodobacteraceae* family and one potentially novel family in the *Cytophagales* order. Functional genomics revealed presumed symbiosis factors typical of specific taxonomic groups (i.e. taurine metabolism genes among the *Alphaproteobacteria,* chitinase encoding genes and eukaryotic-like proteins in the *Cytophagia* genome) while multidrug efflux pumps, also important in host-microbe interactions, were common across all genomes. Moreover, we detected 76 secondary-metabolite biosynthetic gene clusters putatively involved in the production of drug-like compounds or signalling molecules across all genomes, warranting future biotechnologically driven research into their coding potential.

## 1. Introduction

Marine sponges possess a complex and distinct microbial consortium considered to be involved in the provision of nutrients, chemical defence and immunity to their hosts [1]. Although marine sponges have been the source of novel bacterial species [1, 2], yet we lack clear understanding of the diversity, genome structure and metabolic potential of most sponge associates. Apart from the ecological importance and roles of sponges and its microbial community in ecosystem functioning [3], it is now known that the marine sponge microbiome ranks as one of the most prolific sources of bioactive secondary metabolites in the seas [4, 5]. Isolated bacteria permit the full study of the physiology and function of microorganisms [6]. To date, limited information is available for understanding the physiology and function of sponge symbionts and their natural product biosynthesis potential. Although recent studies approaching the coding potential of metagenome-assembled genomes (MAGs) have provided useful insights into symbiosis factors that govern sponge-prokaryote interactions [7, 8], examination of cultivated sponge associated bacteria and their genomes is still much required to address the abovementioned knowledge gaps, especially in regards with the laboratory validation of the observed genomic traits.

In the present study, we describe the general genomic features and provide brief functional information of 14 bacterial strains isolated from the marine sponge *Spongia* sp. in the northeast Atlantic. This study releases new data for the further exploration of complex sponge-microbe interactions and promotes comparative genomics studies of symbiotic microorganisms by adding up novel genome resources to public databases, including original genomes of three novel bacterial lineages unclassifiable at the genus level, the first genome sequence for the genus *Lentilitoribacter* and further genome assemblies of thus-far poorly-studied genera such as *Agarivorans, Sphingorhabdus* and *Tropicibacter*.

## 2. Data description

### 2.1. Sampling, Isolation, and Genome sequencing

In April 2015, *Spongia* sp. specimens (4 biological replicates) were collected by SCUBA diving at 20 m open water stations in Algarve, Faro (Portugal). In the lab, microbial cell suspensions (MCSs) were prepared from sponge samples through grinding and homogenization in artificial seawater as described previously [9]. MCSs were then serially diluted (10^-3^ to 10^-8^) and 100 μL of each dilution were spread plated in triplicates on 1/10R2A [10] and VXA [11] media. Plates were incubated at 18°C for up to 8 weeks. Bacterial growth was monitored weekly and colony forming units (CFUs) were counted. Colonies were selected randomly considering morphologically different ones as well, and pure cultures obtained by streaking colonies on the same isolation media.

In total, 252 colonies (160 from 1/10-R2A and 92 from VXA, Table S1) were selected. Genomic DNA was extracted from cell pellets derived from freshly grown cultures using the Wizard^®^ Genomic DNA Purification Kit (Promega, Madison, USA) according to the manufacturer’s instructions. Genomic DNA samples of all isolates were then subjected to 16S rRNA gene amplification by PCR followed by Sanger sequencing according to established procedures [12]. Taxonomic assignment of bacterial isolates to the genus level was performed using the classifier tool of the ribosomal database project (RDP, release 11, [13]. Identification of closest matches to all sequence queries were also obtained using the BLAST algorithm (February 2018, Table S1).

*Alphaproteobacteria* and *Gammaproteobacteria* strains were frequently isolated from R2A and VXA, respectively (Figure 1). Based on the taxonomic assignments obtained with RDP (Table S2), 14 strains originating from both media were chosen for genome analysis-which in most cases would likely correspond to novel genus/species from sponges (Table S1&S2). These were classified into 11 formally described and 3 potentially novel genera from *Actinobacteria* (3 genera), *Bacteroidetes* (1 putatively new genus and family), and *Proteobacteria* (10 genera from Alpha - 6 strains, with two putatively new genera- and from Gamma - 4 strains - classes) phyla (Table 1).

**Figure 1.**
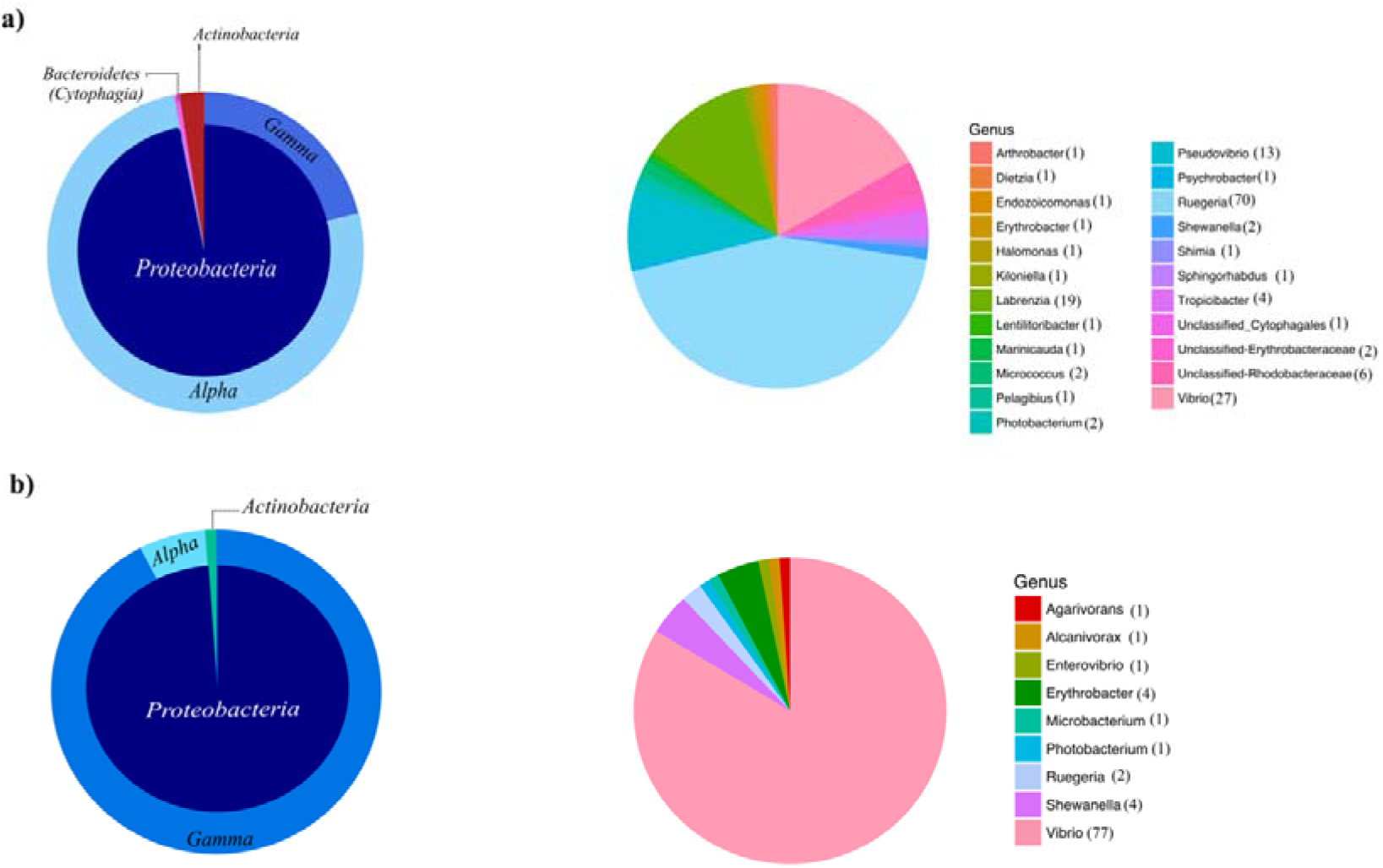
Taxonomic distribution of bacteria retrieved from *Spongia* sp. on **a)** 1/10 R2A **b)** VXA media. The number in brackets indicate the number of isolates obtained for each genus.

**Table 1.**
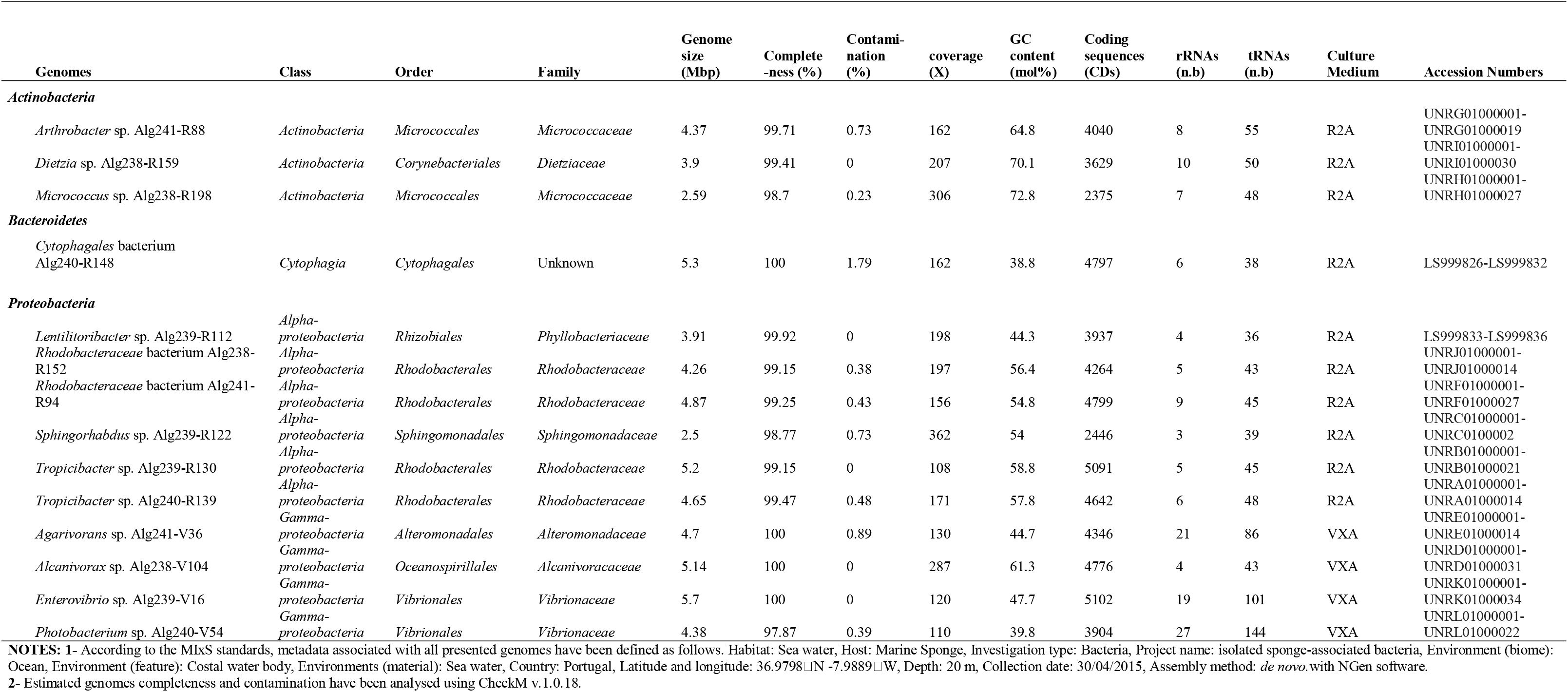
Genome features of all bacterial strains described in this study.

Afterwards, Illumina MiSeq sequencing was carried out at MR DNA (CA, United States). Paired-end libraries (2×301bp) were accomplished and *de novo* assembly was performed with the NGen DNA assembly software by DNAStar, Inc. as described previously [14]. Additionally, CheckM v.1.0.18 was used to assess the completeness and contamination of the generated genomes [15]. Genome annotations were conducted using RAST [16], also using COG database via WebMGA [17]. Furthermore, the genomes were analyzed for the presence of gene clusters indicative of secondary metabolism using antiSMASH version 5.0.1 [18]. Default parameters were used for all software.

### 2.2. Basic Genome Features

The genomic feature information of all strains is outlined in Table 1. All genomes possessed > 99% completeness (except three genomes) and <1% contamination, suggesting high data reliability for further comparative genomic studies, for example.

Three genera including *Arthrobacter* sp. Alg241-R88, *Dietzia* sp. Alg238-R159, *Micrococcus* sp. Alg238-R198 isolated on 1/10R2A were sequenced from phylum *Actinobacteria.* The size of the genomes ranged from 2.59 to 4.37 Mbp with GC content ranging from 64.8 to 72.8 %. Gene annotation using RAST identified 4040, 3629, and 2375 protein-coding sequences (CDs) for strains Alg241-R88, Alg238-R159, and Alg238-R198, respectively. From COG annotations, the genes for glycosyltransferase, acetyltransferases, ABC-type multidrug transport system were assigned for these three strains (Table S3).

Besides, a novel genome from the phylum *Bacteroides* was sequenced. The genome of *Cytophagales* bacterium Alg240-R148 is 5.3 Mbp in length with an overall GC content of 38.8%. Gene annotation using RAST identified a total of 4797 coding sequences, 38 tRNAs, and 6 rRNAs. This genome is 100% complete based on CheckM estimations. The COG annotation indicated genes involved in ABC-type antimicrobial peptide transport system, ABC-type multidrug transport system, chitinases, glycosidases and tetratricopeptide repeats (Table S3).

Moreover, six strains of the class *Alphaptoteobacteria* belonging to the families *Phyllobacteriaceae, Rhodobacteraceae*, and *Sphingomonadaceae* were sequenced. The size of the genomes ranged from 2.5 to 4.78 Mbp with GC content ranging from 44.3 to 58.8%. RAST annotation indicated a range of 2446 coding sequences for *Sphingorhabdus* sp. Alg239-R122 to 5091 for *Tropicibacter* sp. Alg239-R130. In agreement with recent findings on sponge-prokaryote symbiosis factors characteristic of *Alphaproteobacteria* species [15], genes for Glutathione S-transferase, taurine catabolism dioxygenase (except for strain Alg239-R122), thiamine monophosphate synthase, Catalase (peroxidase I) were predicted for all presented *Alphaproteobacteria* strains based on COG annotation.

From the *Gammaproteobacteria* strains obtained on VXA medium, *Agarivorans* sp. Alg241-V36, *Alcanivorax* sp. Alg238-V104, *Enterovibrio* sp. Alg239-V16, and *Photobacterium* sp. Alg240-V54 were sequenced. The first three strains were 100% complete based on CheckM estimations. Genome sizes across the four genera ranged from 4.38 to 5.7 Mbp with GC content ranging from 39.8 to 61.3%. RAST annotation indicated a range of 3904 coding sequences for *Photobacterium* to 5102 for *Enterovibrio.* COG annotations revealed a range of genes involved in multidrug resistance and transport, and coding for collagenase and related proteases, putative threonine efflux proteins and Methyl-accepting chemotaxis proteins for all four strains (Table S3).

### 2.3. Genome-wide secondary metabolite profiling

Totals of 16 (*Actinobacteria*), 3 (*Cytophagia*/*Bacteroidetes*), 38 (*Alphaptoteobacteria*), and 19 (*Gammaproteobacteria*) secondary metabolite biosynthetic gene clusters (BGCs) were detected using antiSMASH (Figure 2). For *Actinobacteria* strains, PKS (Polyketide synthases), siderophore, NRPS (Nonribosomal peptide synthetases), betalactone, terpene and ectoine BGCs were found. For the unclassified *Cytophagales* strain, one bacteriocin, terpene, and arylpolyene BGC each were detected. For all *Alphaproteobacteria* strains, PKS, bacteriocin, NRPS, terpene, homoserine lactone (hserlaton), arylpolyene, betalacton, ectoine, and lassopeptide BGCs were predicted. These findings match those observed in earlier studies [19], particularly the presence of bacteriocin in *Alphaproteobacteria* species. For all *Gammaproteobacteria* strains, shared bacteriocin, betalactone and arylpolyene BGCs were found, though for *Enterovibrio* sp. Alg239-V16 eight different BGCs were predicted (Figure 2). These results highlight the genomic potential of the isolated bacteria for natural product discovery.

**Figure 2.**
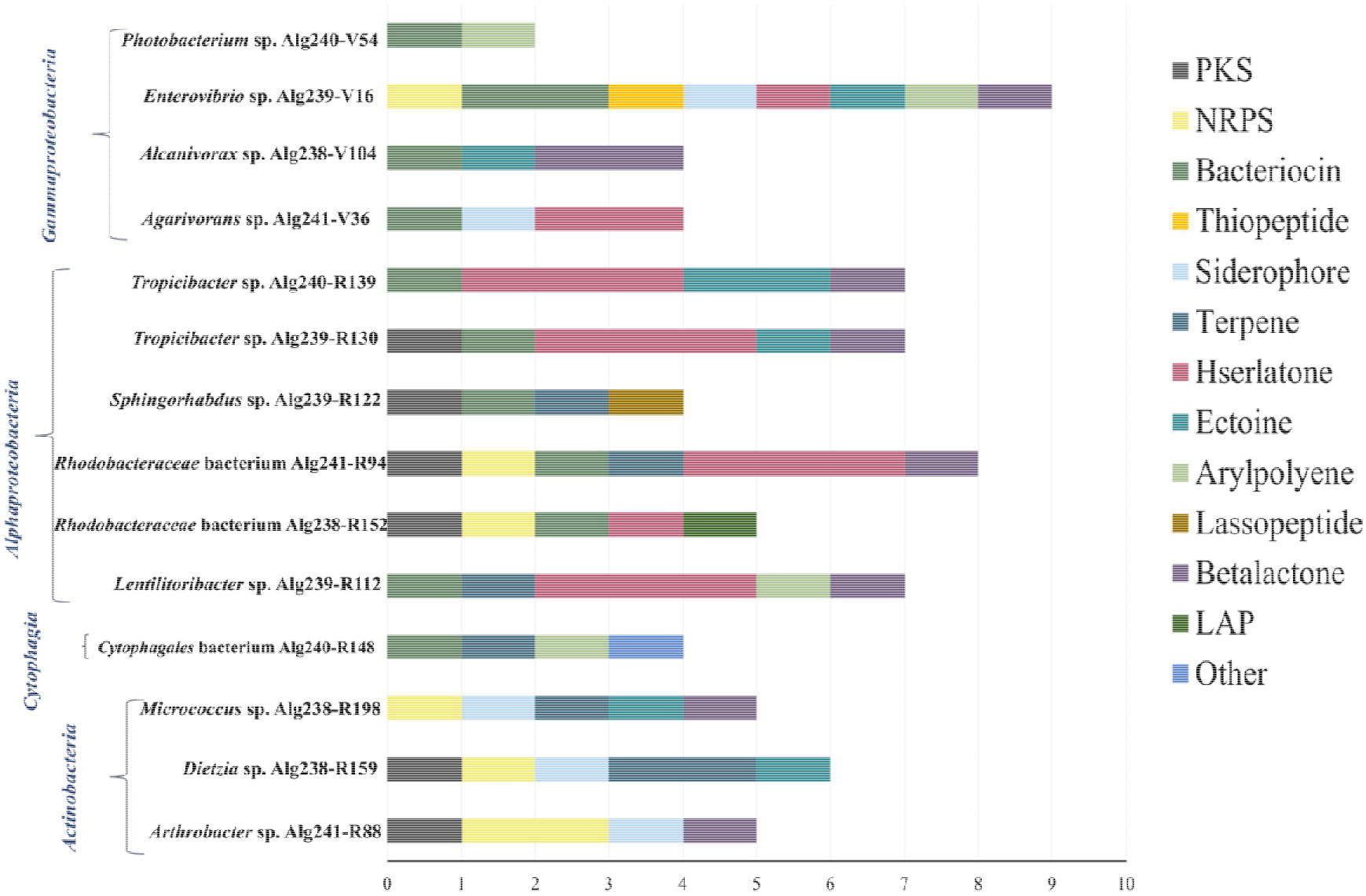
Secondary metabolite biosynthetic gene clusters predicted with antiSMASH across all the studied genomes.

#### Genome sequence accession number

The assembled genome sequences are available in the European Nucleotide Archive - European Molecular Biology Laboratory-EBI under the project number PRJEB28331 and can be accessed using the accession numbers given in Table 1. (http://www.ebi.ac.uk/ena/data/view/ <ACCESSION NUMBER>). Also, 16S rRNA gene sequences of the bacterial isolates (OTUs) were deposited at NCBI GenBank under the accession numbers MH818465 to MH818518 (Table S2).

## Acknowledgments

The authors would like to thank Jorge M.S. Gonçalves’ research team at CCMAR-UAlg for sampling the sponges from the sea. We thank Telma Franco for her help during sample processing and Marta Valente for Sanger-sequencing of 16S rRNA genes. EK would like sincerely to thank Alireza Asvadi for his encouragement to write up the study. This work was performed when EK was PhD student supported by a full scholarship from the Erasmus Mundus Programme/SALAM EMA2 lot7/SALA1206422. Material and resources for the lab experiments were funded by FCT-Fundação para a Ciência e a Tecnologia (Portugal) through the project grant PTDC/BIA-MIC/3865/2012, conceded to RC. The funders had no role in study design, data collection and interpretation, or the decision to submit the work for publication.

## Author contributions

**Elham Karimi:** conceived the study, performed experiments, curated and analysed the data, wrote the original manuscript draft. **Rodrigo Costa**: conceived the study, provided materials, reviewed and edited the original manuscript draft. Both authors reviewed and edited the final manuscript draft.

## Ethics statement

This study relied on *in situ* sampling of microorganisms from marine invertebrates without a nervous system, and as such was exempt from ethical approval procedures according to the current Portuguese legislation (Decreto-Lei n° 113/2013). This study did not occur within privately owned or protected areas. This study did not involve endangered or protected species. The sampling methodology privileged minimally invasive handling procedures, following the guidelines of the European Directive 2010/63/EU.

## Supplementary Material

**Table S1.** 16S rRNA gene-based classification (genus level) and closest 16S rRNA gene relatives of bacterial isolates obtained from *Spongia* sp. on 1/10R2A and VXA media.

**Table S2.** Hierarchical, 16S rRNA gene-based taxonomic affiliation of bacterial isolates retrieved from *Spongia* sp. on **a)** 1/10R2A and **b)** VXA media, with their respective grouping into OTUs at 98% 16S rRNA gene identity.

**Table S3.** COG annotation of sponge-associated bacterial genomes presented in this study.

